# Amyloid Polymorphism of Lysozyme Governs Cross-Seeding of Insulin Aggregation

**DOI:** 10.64898/2026.08.26.747312

**Authors:** Sanjay K. Metkar, Vijay Eerati, Ayyalusamy Ramamoorthy

## Abstract

Amyloid fibrils are highly ordered protein aggregates characterized by a conserved cross-β-sheet architecture despite originating from structurally diverse precursor proteins. Growing evidence suggests that interactions between different amyloidogenic proteins can modulate aggregation pathways through heterologous cross-seeding; however, the influence of seed polymorphism on the structure and biological properties of cross-seeded fibrils remains poorly understood. Here, we investigated the cross-seeding of native human insulin by two structurally distinct polymorphs of hen egg-white lysozyme (HEWL): flexible fibrils (FFs) and rigid fibrils (RFs). Native insulin remained stable under physiological conditions and underwent spontaneous fibrillation only under acidic conditions. In contrast, both HEWL polymorphs efficiently induced insulin aggregation at physiological pH, bypassing the nucleation barrier. Thioflavin T fluorescence, circular dichroism spectroscopy, and transmission electron microscopy revealed that lysozyme FFs templated the formation of insulin flexible fibrils (IFFs), whereas lysozyme RFs produced insulin rigid fibrils (IRFs), demonstrating that the structural characteristics of the parental HEWL polymorphs were propagated during heterologous cross-seeding. The toxicity of the resulting insulin fibrils was evaluated in SH-SY5Y neuronal cells and CCF-STTG1 astrocytes. IFFs exhibited minimal cytotoxicity and only subtle morphological alterations, whereas IRFs caused modest reductions in cell viability accompanied by more pronounced cellular damage. These findings demonstrate that the structural polymorphism of HEWL fibrils governs both the architecture and biological activity of cross-seeded insulin fibrils, highlighting amyloid polymorphism as an important determinant of heterologous amyloid propagation and a potential design principle for engineering functional amyloid-based biomaterials and protein delivery platforms.

## Introduction

Protein misfolding into amyloid fibrils represents a common pathological feature of more than fifty human disorders, including Alzheimer’s disease, Parkinson’s disease, systemic amyloidosis, and type II diabetes mellitus ^1^. These fibrils are distinguished by their highly ordered cross-β architecture^2^. Although traditionally regarded as irreversible pathological end-products of protein misfolding, amyloids are now recognized as structurally and functionally diverse supramolecular assemblies. Beyond their pathological roles, functional amyloids participate in numerous physiological processes, while engineered amyloids have emerged as promising biomaterials for biomedical and nanotechnological applications^3^. This expanding view has shifted attention from amyloid formation itself to the structural diversity, or amyloid polymorphism, that governs their physicochemical properties and biological functions.

Amyloid polymorphism enables a single protein sequence to assemble into multiple fibrillar conformations that differ in molecular packing, morphology, mechanical properties, stability, and biological activity. Recent studies have shown that hen egg-white lysozyme (HEWL) can form two distinct polymorphic states: flexible fibrils (FFs) with a partially ordered, thermally reversible architecture and rigid fibrils (RFs) possessing a highly ordered, irreversible amyloid core^4, 5^. These findings challenged the conventional view that the reversible or irreversible nature of amyloids is solely encoded by the primary amino acid sequence and instead demonstrated that protein folding pathways critically determine amyloid structure and stability^5^. In parallel, increasing evidence has established heterologous amyloid cross-seeding as an important mechanism by which fibrils formed from one protein accelerate the aggregation of another structurally unrelated protein^6^. Unlike homotypic seeding, which occurs between identical proteins, heterologous cross-seeding is primarily driven by structural complementarity of the cross-β surface through hydrogen bonding, hydrophobic interactions, and electrostatic contacts rather than primary sequence similarity^7, 8^. Such interactions enable preformed fibrils to bypass the energetically unfavorable primary nucleation step, propagate conformational information to newly formed fibrils, and generate distinct amyloid strains with unique structural and biological characteristics^9^.

Human insulin provides an ideal model for investigating heterologous amyloid propagation because its fibrillation pathway is well characterized. Under physiological conditions, insulin remains predominantly in its native α-helical conformation and does not readily aggregate unless exposed to destabilizing conditions such as acidic pH, elevated temperature, or prolonged agitation^10, 11^. Consequently, insulin offers a sensitive system for determining whether preformed heterologous amyloid seeds can overcome the kinetic barrier to aggregation and direct fibril formation under non-amyloidogenic conditions. Although hereditary lysozyme amyloidosis is a rare systemic amyloid disease and insulin-derived amyloidosis is typically associated with repeated subcutaneous insulin administration, both proteins share the fundamental propensity to undergo conformational conversion into cross-β-sheet-rich amyloid fibrils^12–14^. The coexistence of multiple amyloidogenic proteins in biological systems has led to increasing recognition that heterologous amyloid interactions may influence disease progression through cross-seeding mechanisms. While a direct pathological interaction between lysozyme and insulin has not been demonstrated *in vivo*, lysozyme amyloid fibrils provide a well-defined model for investigating how the structural properties of one amyloid species modulate the aggregation pathway of an unrelated protein. Such studies offer mechanistic insight into the broader phenomenon of amyloid crosstalk, where pre-existing fibrillar assemblies may alter aggregation kinetics, fibril architecture, and the biological properties of secondary amyloids. Understanding these interactions is particularly relevant because mixed amyloid deposits and amyloid polymorphism are increasingly implicated in the heterogeneity, progression, and tissue-specific manifestations of protein misfolding disorders. Moreover, the structural polymorphism of amyloid fibrils has emerged as a critical determinant of their seeding efficiency, conformational templating, cellular interactions, and pathological outcomes. Distinct polymorphic states may therefore differentially direct the aggregation of heterologous proteins, giving rise to secondary amyloids with unique structural and biological characteristics. Beyond their pathological relevance, certain amyloid polymorphs—particularly structurally flexible assemblies—have attracted interest as functional biomaterials owing to their tunable mechanical properties and potential biocompatibility, whereas highly ordered rigid fibrils provide valuable models for understanding disease-associated amyloid conformations. Consequently, elucidating how structurally distinct HEWL polymorphs (HEWL-FFs and HEWL-RFs) influence insulin aggregation offers not only mechanistic insight into amyloid cross-seeding but also a framework for exploring the dual roles of amyloid polymorphism in pathological protein aggregation and the rational design of amyloid-based functional materials.

Recently, HEWL amyloid fibrils were shown to efficiently cross-seed insulin aggregation through structure-based molecular recognition involving stable non-covalent interactions, establishing lysozyme as an effective heterologous nucleating agent for insulin fibrillation. However, these studies did not address whether distinct polymorphic forms of HEWL fibrils produce structurally and biologically different insulin amyloid polymorphs^15^. Here, we investigated how the polymorphic state of HEWL fibrils influences the heterologous cross-seeding of native human insulin. We first established that native insulin remains structurally stable under physiological conditions and undergoes amyloid fibrillation only under destabilizing conditions, highlighting the requirement for an external nucleating trigger. We then demonstrate that both HEWL -FFs and HEWL-RFs efficiently overcome this kinetic barrier and template insulin aggregation under physiological conditions. Remarkably, HEWL-FFs generate flexible insulin fibrils (IFFs) that exhibit minimal cytotoxicity toward neuronal and astrocytic cells, whereas HEWL-RFs produce rigid insulin fibrils (IRFs) that showed cytotoxicity. Collectively, our findings reveal that the polymorphic state of the heterologous seed governs not only the structural phenotype of the resulting insulin fibrils but also their biological activity. These results provide new mechanistic insight into amyloid polymorphism during heterologous cross-seeding and establish fibril architecture as a key determinant of amyloid function, with potential implications for both amyloid-associated diseases and the rational design of functional amyloid-based biomaterials.

## Results and Discussion

### Structural Characterization of Native Insulin and HEWL Amyloid Polymorphs

To establish the experimental basis for heterologous cross-seeding, the structural and aggregation properties of native insulin and HEWL amyloid polymorphs were first characterized using Thioflavin T (ThT) fluorescence, far-UV circular dichroism (CD) spectroscopy, and transmission electron microscopy (TEM) (Figure 1). Native human insulin is conformationally stable under physiological conditions and generally requires destabilizing environments, such as acidic pH and elevated temperature, to undergo amyloid fibrillation. Accordingly, insulin (100 μM) incubated under acidic conditions (pH 3) exhibited the characteristic sigmoidal ThT fluorescence profile consisting of a lag phase followed by rapid fibril growth and a plateau phase, indicative of nucleation-dependent amyloid formation (Figure 1a). Structural conversion was confirmed by CD spectroscopy, where native insulin displayed the characteristic α-helical spectrum with minima near 208 and 222 nm, whereas insulin amyloid exhibited a dominant minimum around 218 nm, consistent with a β-sheet-rich amyloid conformation^16^ (Figure 1b). TEM further confirmed the formation of mature insulin amyloid fibrils as long, unbranched filamentous structures characteristic of cross-β assemblies (Figure 1c). These observations demonstrate that insulin readily undergoes fibrillation only under amyloidogenic conditions, whereas under physiological conditions it remains predominantly in its native conformation, providing an appropriate negative control for subsequent cross-seeding experiments.

**Figure 1.**
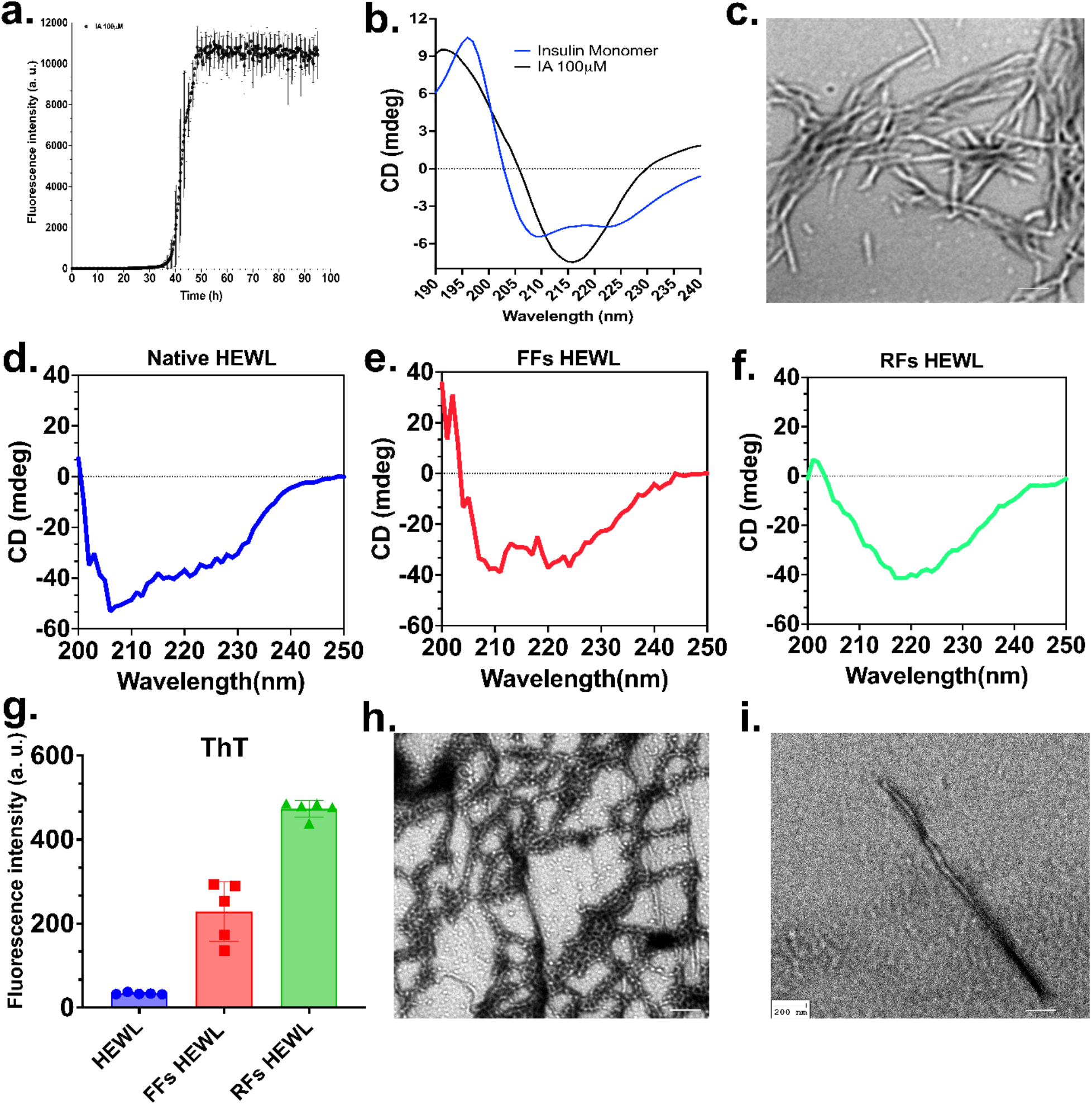
Native human insulin undergoes spontaneous amyloid fibrillation under acidic conditions. (a) ThT fluorescence kinetics of 100 μM human insulin incubated under acidic conditions (pH 3) demonstrating spontaneous amyloid fibril formation after a characteristic nucleation phase followed by rapid fibril growth. (b) Far-UV circular dichroism (CD) spectra of native insulin monomer and insulin fibrils formed under acidic conditions, showing the transition from the native α-helical conformation to a β-sheet-rich amyloid structure. (c) Representative TEM image of mature insulin amyloid fibrils formed under acidic conditions, exhibiting long, unbranched fibrillar morphology characteristic of cross-β amyloid assemblies. (d–f) Far-UV CD spectra of native HEWL, FFs HEWL, and RFs HEWL, respectively, showing distinct secondary structural conformations. (g) End-point ThT fluorescence intensity of native and fibril forms of HEWL confirming amyloid formation with higher ThT binding by RFs than FFs. (h, i) Representative TEM images of FFs HEWL and RFs HEWL, illustrating their distinct fibrillar morphologies. Data in (g) are presented as mean ± SD from independent experiments. Scale bar = 200 nm.

The structural characteristics of native HEWL and its two polymorphic amyloid forms, HEWL-FFs and HEWL-RFs, were subsequently examined (Figure 1d–i). Far-UV CD spectroscopy revealed a pronounced conformational transition accompanying amyloid formation. Native HEWL exhibited the characteristic spectrum of an α-helical protein (Figure 1d), whereas both HEWL-FFs and HEWL-RFs displayed spectra dominated by a negative band centered near 218 nm, indicative of β-sheet-rich amyloid structures (Figure 1e, f). Although both polymorphs adopted the canonical cross-β conformation, the HEWL-RFs spectrum exhibited a deeper negative ellipticity around 218 nm than HEWL-FFs, suggesting a more ordered and extensively packed β-sheet core.

These structural differences were further corroborated by ThT fluorescence measurements (Figure 1g). Native HEWL exhibited negligible fluorescence, whereas HEWL-FFs showed a marked increase in ThT intensity, confirming amyloid formation. HEWL-RFs produced the highest fluorescence signal, consistent with greater β-sheet stacking and enhanced fibrillar order. TEM analysis further distinguished the two polymorphs (Figure 1h,i). HEWL-FFs formed short, curved, and highly interconnected fibrillar networks composed of loosely packed filaments, whereas HEWL-RFs appeared as long, straight, densely packed needle-like fibrils characteristic of highly ordered amyloid assemblies.

To exclude potential contributions from HEWL monomers or preformed fibril seeds to the ThT fluorescence observed during cross-seeding, control experiments were performed using native insulin incubated with monomeric HEWL and HEWL fibril seeds alone (Supplementary Figure S1). Native insulin (100 μM) incubated with increasing concentrations of monomeric HEWL (2.5–10 μM) at physiological pH exhibited no appreciable increase in ThT fluorescence throughout the incubation period, confirming that monomeric HEWL does not induce spontaneous insulin amyloid formation under these conditions (Supplementary Figure S1a). In parallel, HEWL-FFs and HEWL-RFs seeds alone produced only low, stable ThT fluorescence signals over the course of the experiment (Supplementary Figure S1b, c), consistent with the intrinsic ThT binding of preformed HEWL fibrils. These control experiments demonstrate that the background ThT signal arising from the HEWL seeds is minimal and remains essentially unchanged during the assay. Therefore, increase in ThT fluorescence observed in the cross-seeding experiments predominantly reflects the formation of newly generated insulin amyloid fibrils rather than fluorescence originating from the seed material itself. Collectively, these findings validate that the enhanced ThT responses and the differences in aggregation kinetics reported for HEWL-FFs and HEWL-RFs mediated cross-seeding are attributable to polymorph-dependent templated insulin fibrillization rather than intrinsic fluorescence contributions from HEWL monomers or preformed fibril seeds.

Collectively, these findings demonstrate that although both HEWL-FFs and HEWL-RFs possess the canonical cross-β amyloid architecture, they exhibit distinct differences in secondary structure, β-sheet organization, ThT binding, and supramolecular morphology. Importantly, the stability of native insulin under physiological conditions, together with the successful generation of two structurally distinct HEWL amyloid polymorphs, establishes an experimental platform for investigating how the conformational state of a heterologous amyloid seed influences insulin aggregation, the structural characteristics of the resulting fibrils, and their biological activity. Accordingly, native insulin was incubated with increasing concentrations of HEWL-FFs or HEWL-RFs under physiological conditions to evaluate their heterologous cross-seeding capability. The resulting insulin fibrils were subsequently characterized by ThT fluorescence, CD spectroscopy, TEM, and cellular assays to determine how HEWL amyloid polymorphism governs the formation, architecture, and biological properties of cross-seeded insulin fibrils.

### FFs of HEWL fibrils efficiently cross-seed insulin into flexible amyloid fibrils with minimal cytotoxicity

To determine whether HEWL-FFs template the formation of insulin flexible fibrils (IFFs), native human insulin (100 μM) was incubated with increasing concentrations of HEWL-FFs (2.5, 5, and 10 μM). ThT fluorescence revealed a concentration-dependent increase in amyloid formation compared with insulin alone (Figure 2a), indicating efficient cross-seeding by HEWL-FFs. The seeded samples exhibited accelerated fibril formation with a markedly reduced lag phase, demonstrating that preformed HEWL-FFs effectively bypass the nucleation barrier and template insulin aggregation. These findings are consistent with the concept that heterologous amyloid cross-seeding proceeds through secondary nucleation on the surface of pre-existing fibrils, thereby eliminating the need for spontaneous nucleus formation^6^.

**Figure 2.**
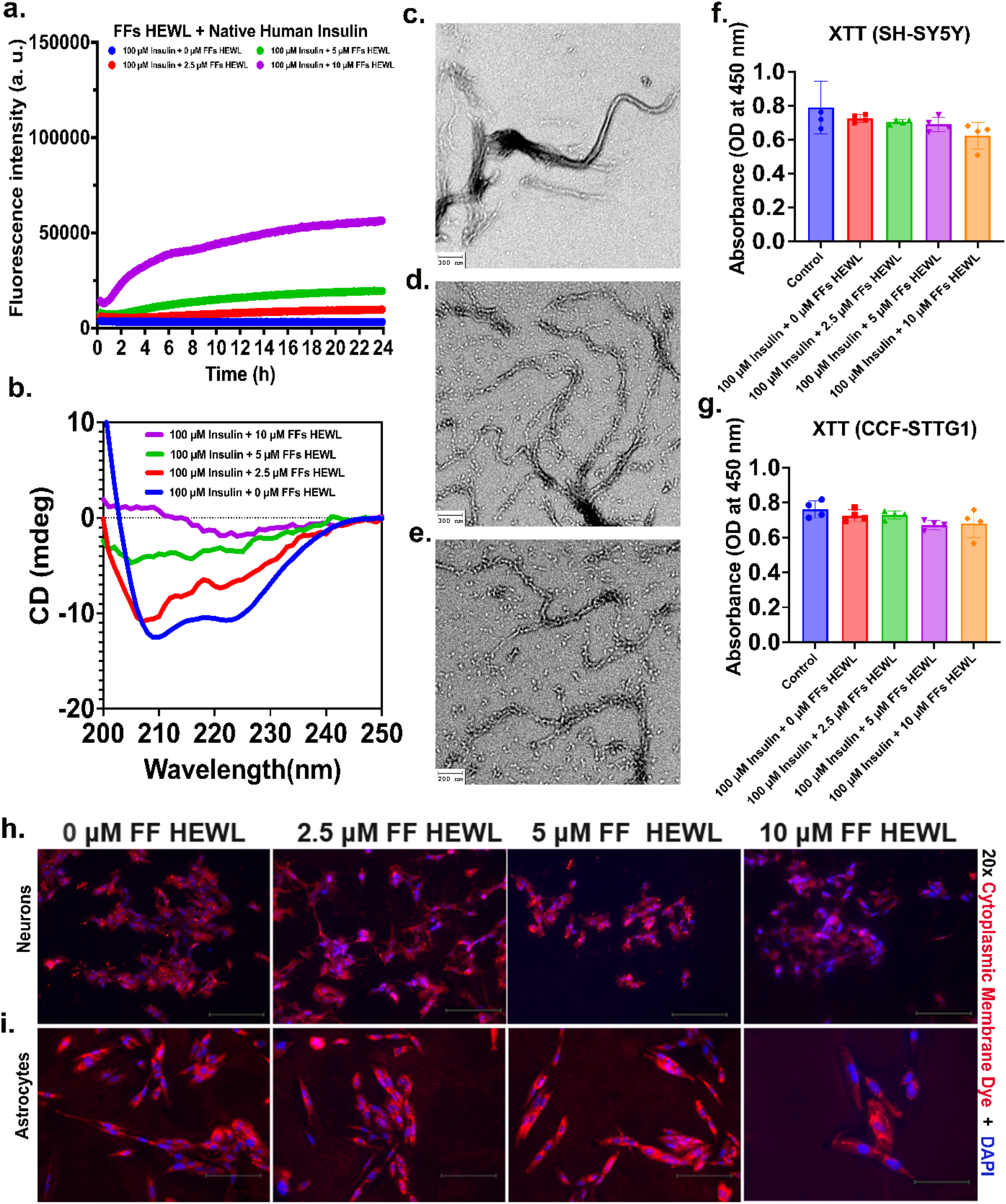
FFs of HEWL cross-seed the formation of insulin flexible fibrils (IFFs) and their interactions with central nervous system (CNS) cell lines. (a) Thioflavin T (ThT) fluorescence kinetics of 100 μM native human insulin incubated alone or with increasing concentrations of FFs of HEWL (2.5, 5, or 10 μM) for 24 h, demonstrating concentration-dependent cross-seeding and the formation of IFFs. (b) Far-UV circular dichroism (CD) spectra showing the conversion of native α-helical insulin to a β-sheet-rich conformation following cross-seeding by FFs of HEWL. (c–e) Representative transmission electron microscopy (TEM) images of insulin FFs generated by cross-seeding with FFs of HEWL (scale bars: 300, 300, and 200 nm, respectively). (f, g) XTT cell viability of SH-SY5Y neuronal cells (f) and CCF-STTG1 astrocyte cells (g) following treatment with native insulin or insulin FFs generated in the presence of FFs of HEWL (2.5, 5, or 10 μM). Data are presented as mean ± SD (n = 4), indicating no significant cytotoxicity under the experimental conditions. (h, i) Representative fluorescence micrographs (20×) of primary neurons (h) and astrocytes (i) treated with the corresponding insulin FF preparations generated by cross-seeding with FFs of HEWL and stained with a plasma membrane dye (red) and DAPI (blue), showing concentration-dependent cellular interactions and morphological changes. Scale bar = 150 μm.

The structural conversion of insulin to IFFs was confirmed by far-UV CD spectroscopy (Figure 2b). Native insulin displayed the characteristic α-helical spectrum with minima at approximately 208 and 222 nm, whereas cross-seeded samples progressively lost their α-helical signature and developed a dominant minimum near 218 nm, consistent with the formation of β-sheet-rich amyloid fibrils. The increase in β-sheet content correlated well with the enhanced ThT fluorescence, confirming that the observed fluorescence increase resulted from amyloid formation.

Further, TEM verified fibril formation (Figure 2c–e). Cross-seeded insulin formed abundant, elongated, unbranched fibrils exhibiting a flexible, curvilinear morphology resembling the parent HEWL- FFs. At higher seed concentrations, fibrils formed dense interconnected networks, supporting efficient propagation of the flexible fibrillar architecture during heterologous cross-seeding. The preservation of the flexible fibrillar morphology further suggests that the supramolecular architecture of the parental HEWL fibrils is transmitted to the newly formed insulin aggregates, highlighting the ability of amyloid polymorphs to template not only fibril formation but also fibril morphology.

Previous computational and biophysical studies have demonstrated that lysozyme fibrils interact strongly with insulin through hydrogen bonding, hydrophobic contacts, and complementary electrostatic interactions, producing stable heterotypic complexes that facilitate conformational conversion. These observations suggest that the efficient cross-seeding observed here is driven primarily by structural complementarity of the cross-β fibril surface rather than primary sequence similarity, allowing insulin monomers to be recruited onto the HEWL fibril template and converted into β-sheet-rich fibrils^15^.

The biological effects of the cross-seeded insulin fibrils were evaluated using SH-SY5Y neuronal cells and CCF-STTG1 astrocytes. XTT assays showed no significant reduction in cell viability following treatment with insulin flexible fibrils (IFFs) generated in the presence of HEWL-FFS (Figure 2f,g), indicating that these fibrillar assemblies exhibit minimal acute cytotoxicity under the experimental conditions. Fluorescence microscopy of neurons and astrocytes (Figure 2h,i) stained with a plasma membrane marker and DAPI revealed concentration-dependent interactions between the fibrils and CNS cells, with only subtle morphological changes observed at higher fibril concentrations (20× objective; scale bar = 150 μm). No evidence of extensive membrane disruption or widespread cell loss was detected, consistent with the XTT results.

Interestingly, despite their efficient cross-seeding capability, the IFFs elicited only minimal cellular responses. This observation indicates that efficient amyloid propagation does not necessarily correlate with cytotoxicity and supports the emerging view that the biological activity of amyloid assemblies is strongly influenced by their structural polymorphism. The relatively low cytotoxicity of the FFs suggests that their dynamic and less densely packed architecture may interact differently with cellular membranes than highly ordered rigid fibrils.

Collectively, these findings demonstrate that HEWL-FFs efficiently cross-seed native human insulin, promoting its conversion from an α-helical monomer into flexible amyloid fibrils. The resulting fibrils retain the flexible morphology of the parental HEWL seeds while exhibiting minimal toxicity toward neuronal and astrocytic cells. Together with previous mechanistic studies of lysozyme-mediated cross-seeding, these results support a model in which structure-based molecular recognition, rather than sequence homology, governs heterologous amyloid propagation. Furthermore, our data reveal that the structural polymorphism of the seed influences not only the morphology of the resulting insulin fibrils but also their biological response, emphasizing that amyloid architecture is a key determinant of cross-seeding outcomes. If the reversible conversion of these IFFs to native insulin monomers is confirmed experimentally, such assemblies may provide a basis for developing reversible amyloid-based protein delivery platforms, including sustained insulin-release systems^17^.

### RFs of HEWL efficiently cross-seed insulin into rigid amyloid fibrils with enhanced cytotoxicity

To determine whether HEWL-RFs can heterologously seed insulin aggregation and generate insulin rigid fibrils (IRFs), native human insulin (100 μM) was incubated with increasing concentrations of HEWL-RFs (2.5, 5, and 10 μM). ThT fluorescence revealed a concentration-dependent increase in amyloid formation compared with insulin alone (Figure 3a), demonstrating efficient cross-seeding by HEWL-RFs. The seeded reactions exhibited accelerated fibril formation with a markedly reduced lag phase, indicating that preformed HEWL-RFs effectively bypass the nucleation barrier and directly template insulin aggregation. This observation is consistent with the mechanism of heterologous amyloid cross-seeding, whereby pre-existing cross-β fibrils recruit soluble monomers onto their surface and promote secondary nucleation, thereby circumventing the energetically unfavorable primary nucleation step.

**Figure 3.**
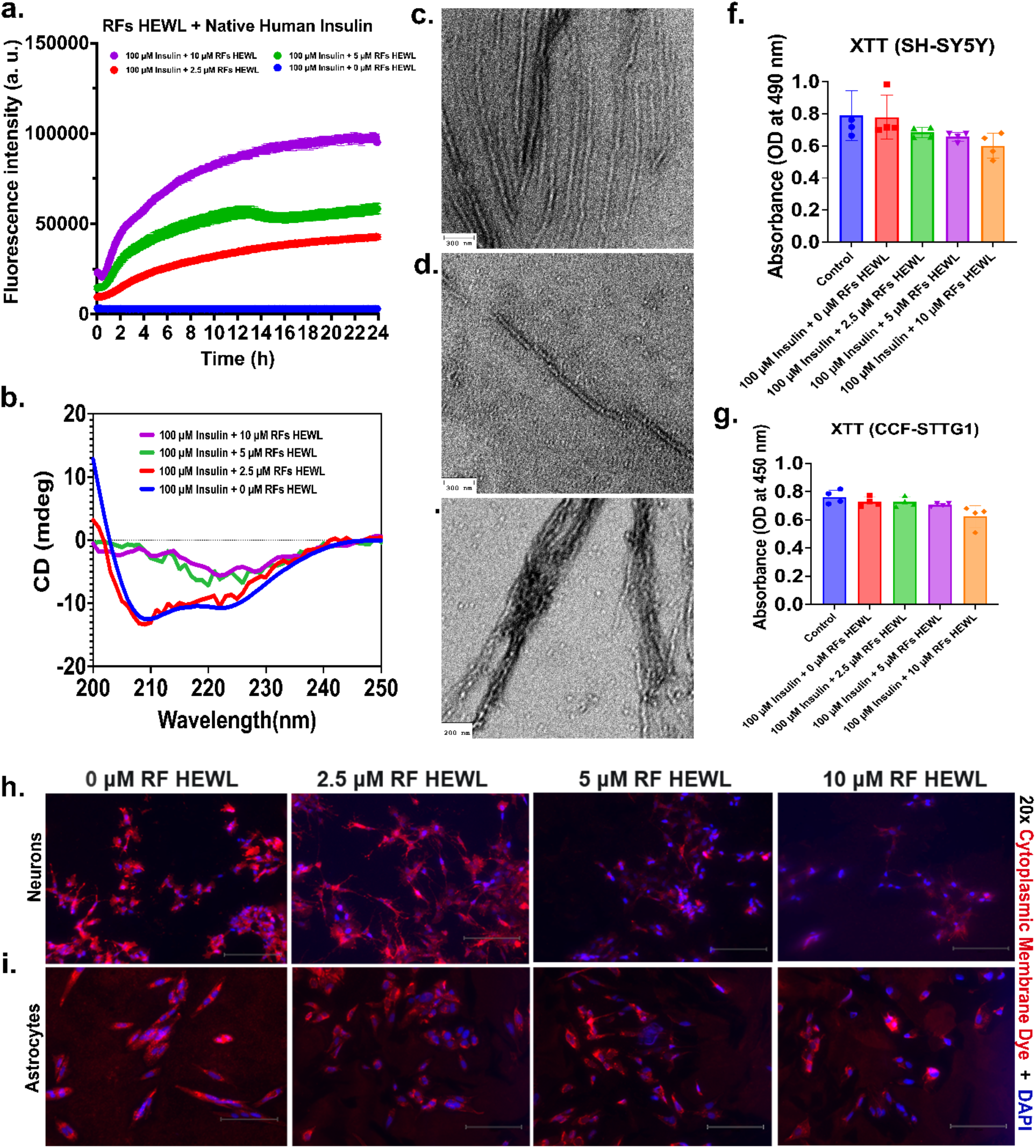
RFs of HEWL cross-seed the formation of insulin rigid fibrils (IRFs) and induce cytotoxic responses in CNS cell lines. (a) ThT fluorescence kinetics of 100 μM native human insulin incubated alone or with increasing concentrations of RFs of HEWL (2.5, 5, or 10 μM) for 24 h, demonstrating concentration-dependent cross-seeding and the formation of IRFs. (b) Far-UV CD spectra showing the conversion of native α-helical insulin into a β-sheet-rich amyloid conformation following cross-seeding by RFs of HEWL. (c–e) Representative TEM images of insulin rigid fibrils generated by cross-seeding with RFs of HEWL (scale bars: 300, 300, and 200 nm, respectively). (f, g) XTT cell viability of SH-SY5Y neuronal cells (f) and CCF-STTG1 astrocytes (g) following treatment with native insulin or IRFs generated in the presence of RFs of HEWL (2.5, 5, or 10 μM). Data are presented as mean ± SD (n = 4). (h, i) Representative fluorescence micrographs (20×) of primary neurons (h) and astrocytes (i) treated with the corresponding IRFs preparations generated by cross-seeding with RFs of HEWL and stained with a plasma membrane dye (red) and DAPI (blue), illustrating concentration-dependent cellular damage and morphological alterations. Scale bar = 150 μm.

Far-UV CD spectroscopy confirmed the structural conversion of insulin following cross-seeding (Figure 3b). Native insulin displayed the characteristic α-helical spectrum with minima at approximately 208 and 222 nm, whereas cross-seeded samples progressively lost their α-helical signature and developed a dominant minimum near 218 nm, confirming the formation of β-sheet-rich amyloid fibrils. The increase in β-sheet content correlated well with the enhanced ThT fluorescence, demonstrating efficient conversion of native insulin into the amyloid state.

TEM further verified fibril formation (Figure 3c–e). Cross-seeded insulin formed abundant, elongated, straight, highly ordered, and densely packed fibrils, characteristic of rigid amyloid assemblies. Extensive lateral association and fibril bundling became increasingly prominent with increasing seed concentration, indicating efficient propagation of the rigid fibrillar architecture. Unlike the IFFs generated by HEWL-FFs, the IRFs faithfully preserved the highly ordered morphology of the parental HEWL-RFs, suggesting that the structural polymorphism of the seed dictates the supramolecular organization of the newly formed IRFs.

The biological consequences of the cross-seeded IRFs were evaluated using SH-SY5Y neuronal cells and CCF-STTG1 astrocytes. XTT assays demonstrated a concentration-dependent reduction in cell viability following treatment with IRFs generated in the presence of HEWL (Figure 3f, g), indicating significantly greater cytotoxicity than that observed for the IFFs. Fluorescence microscopy of primary neurons and astrocytes (20× objective; scale bar = 150 μm) revealed marked morphological alterations characterized by reduced cell density, compromised membrane integrity, and loss of normal cellular morphology, particularly at higher seed concentrations. These observations closely paralleled the XTT measurements and indicate that IRFs interact more aggressively with CNS cells than their IFFs.

The increased cytotoxicity associated with IRFs suggests that fibril architecture is a major determinant of biological activity. The highly ordered and tightly packed structure of rigid fibrils may provide a more stable interface for membrane interaction, facilitating stronger fibril-cell interactions that ultimately impair cellular function. In contrast, the more dynamic architecture of FFs appears to reduce these interactions despite exhibiting comparable cross-seeding efficiency.

Collectively, these findings demonstrate that HEWL-RFs efficiently cross-seed native human insulin, promoting its conversion into β-sheet-rich IRFs that faithfully inherit the morphology of the parental HEWL seeds. Studies showing that HEWL-mediated cross-seeding is governed by structure-based molecular recognition and heterotypic non-covalent interactions, the present results establish that the polymorphic state of the HEWL seed dictates both the supramolecular architecture and the biological activity of the resulting insulin fibrils. The direct comparison between flexible and rigid polymorphs therefore identifies fibril architecture as a critical determinant of heterologous amyloid propagation and cytotoxicity, providing new insight into how distinct amyloid polymorphs may differentially influence amyloid-associated pathology and the rational design of functional amyloid-based biomaterials.

The divergent behavior of HEWL-FFs and HEWL-RFs polymorphs may have broader implications beyond the present model system. HEWL-FFs, which promote structurally adaptable and less cytotoxic insulin assemblies, may be advantageous for the rational design of amyloid-based biomaterials. In contrast, HEWL-RFs generated highly ordered insulin aggregates with enhanced cytotoxicity, suggesting that such polymorphs more closely resemble the pathogenic amyloid conformations implicated in protein misfolding diseases. Although HEWL amyloidosis itself is a rare disorder, these findings provide mechanistic insight into how amyloid polymorphism may govern heterologous cross-seeding events in more clinically relevant diseases.

## Conclusion

This study demonstrates that the polymorphic state of HEWL amyloid fibrils is a critical determinant of heterologous insulin aggregation and the properties of the resulting amyloid assemblies. Although native insulin remained stable under physiological conditions, both HEWL-FFs and HEWL-RFs efficiently bypassed the nucleation barrier and induced insulin fibrillation through heterologous cross-seeding. Importantly, the structural characteristics of the parental HEWL polymorphs were faithfully propagated to the newly formed insulin fibrils, generating IFFs or IRFs with distinct morphologies and biological activities. While IFFs exhibited minimal cytotoxicity toward neuronal and astrocytic cells, IRFs induced cellular damage, demonstrating that seed polymorphism governs not only amyloid architecture but also biological function. These findings establish amyloid polymorphism as a key regulator of heterologous amyloid propagation and provide new mechanistic insight into how structurally distinct amyloid templates dictate the formation and biological behavior of secondary amyloids. The combination of efficient templating, structural flexibility, and low cytotoxicity suggests that IFFs may represent a promising platform for the future development of reversible amyloid-based biomaterials, including long-acting protein depots and sustained-release therapeutic delivery systems. In contrast, the cytotoxicity associated with IRFs provides insight into how pathogenic amyloid polymorphs may differentially contribute to disease progression. Collectively, these findings advance our understanding of amyloid cross-seeding while establishing amyloid polymorphism as a design principle for engineering protein assemblies with tunable structural and functional properties.

## Material and Methods

### Materials

Sodium chloride and sodium phosphate were purchased from Fisher Scientific (Hampton, NH, USA). Chicken egg white lysozyme (HEWL; Cat. No. J60701.14) was purchased from Thermo Fisher Scientific (Waltham, MA, USA) and used without further purification. Thioflavin T (ThT) was obtained from MilliporeSigma (Burlington, MA, USA). Formvar/carbon-coated copper grids (300 mesh) were purchased from MilliporeSigma, and Uranyless stain was obtained from Electron Microscopy Sciences (Hatfield, PA, USA). Recombinant human insulin was obtained from Roche (Indianapolis, IN, USA). SH-SY5Y human neuroblastoma cells and CCF-STTG1 human astrocytoma cells were obtained from the American Type Culture Collection (ATCC, Manassas, VA, USA). Dulbecco’s Modified Eagle Medium (DMEM), RPMI 1640, fetal bovine serum (FBS), penicillin–streptomycin, phosphate-buffered saline (PBS), and 0.25% trypsin–EDTA were purchased from Corning Inc. (Corning, NY, USA). Cell viability was assessed using the CyQUANT™ XTT Cell Viability Assay (Invitrogen™, Thermo Fisher Scientific). BioTracker™ 555 Orange Cytoplasmic Membrane Dye, 4% paraformaldehyde, and Fluoromount-G™ containing DAPI were used for fluorescence imaging. Eight-well chamber slides and sterile 96-well tissue culture plates were used for cell culture experiments.

### Preparation of Flexible and Rigid HEWL Amyloid Fibrils

FFs and RFs of HEWL were prepared following the method reported by Frey et al.,^5^ with minor modifications. Briefly, HEWL (20 mg/mL) was dissolved in 10 mM sodium phosphate buffer (pH 7.0) containing 20 mM tris(2-carboxyethyl)phosphine (TCEP) and 10 mM NaCl. The solution was incubated at 85 °C with continuous shaking at 300 rpm. FFs were prepared by incubating the solution for 5 min, whereas RFs were obtained by extending the incubation to 3 h. Excess TCEP was removed using Amicon Ultra-0.5 centrifugal filter units (10 kDa molecular weight cut-off). The purified reduced HEWL retained in the filter was collected and immediately used for fibril preparation.

### Transmission Electron Microscopy (TEM)

The morphology of FFs and RFs was examined by TEM. Briefly, 10 μL of freshly prepared fibril suspension was placed onto Formvar/carbon-coated 300-mesh copper grids and allowed to adsorb for 10 min at room temperature. Excess sample was removed, and the grids were negatively stained with 10 μL of Uranyless stain for 1 min. After removing excess stain, the grids were air-dried for 10 min before imaging. TEM images were acquired using a Hitachi HT7800 transmission electron microscope operated at an accelerating voltage of 100 kV. Images were collected from at least three independent regions of each grid.

### Thioflavin T (ThT) Fluorescence Assay

Amyloid fibril formation was monitored using the ThT fluorescence assay. HEWL monomers, FFs, or RFs (100 μM) were mixed with 30 μM ThT in 10 mM sodium phosphate buffer (pH 7.0) and transferred to black 384-well microplates. Samples were incubated at 37 °C with orbital shaking at 700 rpm. Fluorescence was measured using a Thermo Scientific Varioskan ALF microplate reader with excitation and emission wavelengths of 440 and 480 nm, respectively. Buffer-only and ThT-only wells served as background controls. Data were background-corrected and averaged from at least three independent experiments.

### Far-UV Circular Dichroism (CD) Spectroscopy

The secondary structures of HEWL monomers, FFs, and RFs were analyzed using a Chirascan circular dichroism spectropolarimeter (Applied Photophysics, UK). Samples were diluted to 50 μM in 10 mM sodium phosphate buffer (pH 7.0), and spectra were recorded from 200 to 250 nm at room temperature using a 1 mm path-length quartz cuvette. Three scans were averaged after baseline subtraction, and the spectra were processed using GraphPad Prism (version 11.0.2).

### Cross-Seeding Aggregation Kinetics by ThT Fluorescence

ThT fluorescence kinetics assays were performed to monitor the cross-seeded aggregation of insulin in the presence of preformed FFs or RFs of HEWL. Fluorescence measurements were acquired using a Biotek Synergy H1 microplate reader with excitation at 452 nm and emission at 485 nm. Briefly, 100 μM native insulin monomer was incubated with 0, 2.5, 5, or 10 μM HEWL FFs or RFs in 10 mM sodium phosphate buffer (pH 7.0) containing 25 μM ThT. Samples were incubated at 37 °C without shaking, and fluorescence was recorded in black 96-well microplates at 15 min intervals for 24 h. Each experiment was independently repeated at least three times to ensure reproducibility. Following completion of the aggregation assay, the resulting insulin aggregates were characterized by CD spectroscopy to evaluate secondary structural changes and by TEM to examine fibril morphology.

### Cell Culture, Cytotoxicity Evaluation, and Fluorescence Microscopy

#### Cell Culture

Human SH-SY5Y neuroblastoma cells were cultured in Dulbecco’s Modified Eagle Medium (DMEM), whereas CCF-STTG1 (CCF) human astrocytoma cells were maintained in Roswell Park Memorial Institute (RPMI-1640) medium. Both media were supplemented with 10% fetal bovine serum (FBS) and 1% penicillin–streptomycin. Cells were maintained at 37 °C in a humidified incubator with 5% CO₂ and passaged upon reaching approximately 70–80% confluency. For routine subculturing, the culture medium was removed, cells were rinsed with phosphate-buffered saline (PBS), detached using 0.25% trypsin–EDTA, collected by centrifugation, resuspended in fresh complete medium, and seeded into new culture vessels for subsequent experiments

#### XTT Cell Viability Assay

Cell viability was assessed using the CyQUANT™ XTT Cell Viability Assay (Cat. No. X12223; Invitrogen™). SH-SY5Y and CCF cells were seeded into 96-well plates at a density of 1 × 10⁴ cells per well and allowed to attach for 24 h before treatment Cells were subsequently treated with 25 μM native insulin monomer, IFFs, IRFs for 24 h. Untreated cells and buffer-treated cells served as negative controls, whereas FFs and RFs were included as seed-only controls. Following incubation, XTT reagent was added directly to each well according to the manufacturer’s protocol, and absorbance was measured at 450 nm using a Thermo Scientific Varioskan ALF microplate reader.

#### Fluorescence Microscopy

For fluorescence imaging, SH-SY5Y and CCF cells were seeded into 8-well chamber slides (CELLTREAT Scientific Products, Cat. No. 50-114-9054) at a density of 1 × 10⁴ cells per well and cultured for 24 h before treatment. Cells were then incubated with native insulin, IFFs, or IRFs for 24 h. Following treatment, living cells were stained with BioTracker™ 555 Orange Cytoplasmic Membrane Dye (SCT107; Sigma-Aldrich) diluted 1:1000 in complete culture medium and incubated for 30 min at 37 °C to visualize plasma membrane integrity. Cells were gently washed with PBS and fixed in 4% paraformaldehyde for 5 min at room temperature. After three additional PBS washes, samples were mounted using Fluoromount-G™ mounting medium containing DAPI (Invitrogen™, Thermo Fisher Scientific) to counterstain nuclei. Slides were stored at 4 °C before imaging. Fluorescence images were acquired using an Invitrogen™ EVOS™ M7000 Imaging System equipped with a 20× objective. Membrane morphology was visualized using the red fluorescence channel, while nuclei were imaged using the DAPI channel. Representative images were collected using identical acquisition settings for all experimental groups to enable qualitative comparison of treatment-induced changes in cellular morphology and nuclear integrity.

## Acknowledgements

We acknowledge the Biological Science Imaging Resource (BSIR) and the Institute of Molecular Biophysics (IMB) for access to core facilities. We thank Peter Randolph (IMB) and Anthony Warrington (BSIR) for their technical support.

## Author Contributions

Sanjay Metkar: Investigation, Methodology, Data curation, Formal analysis, Writing – original draft and review & editing. Vijay Erati : Investigation, Data curation, Writing – review & editing. Ayyalusamy Ramamoorthy: Conceptualization, Supervision, Funding acquisition, Project administration, Writing – review & editing.

## Data Availability

All data supporting the findings of this study are included in the article and its Supporting Information. Raw experimental data are available upon reasonable request.

