## Supporting Information for "Amyloid Polymorphism of Lysozyme Governs Cross-Seeding of Insulin Aggregation"

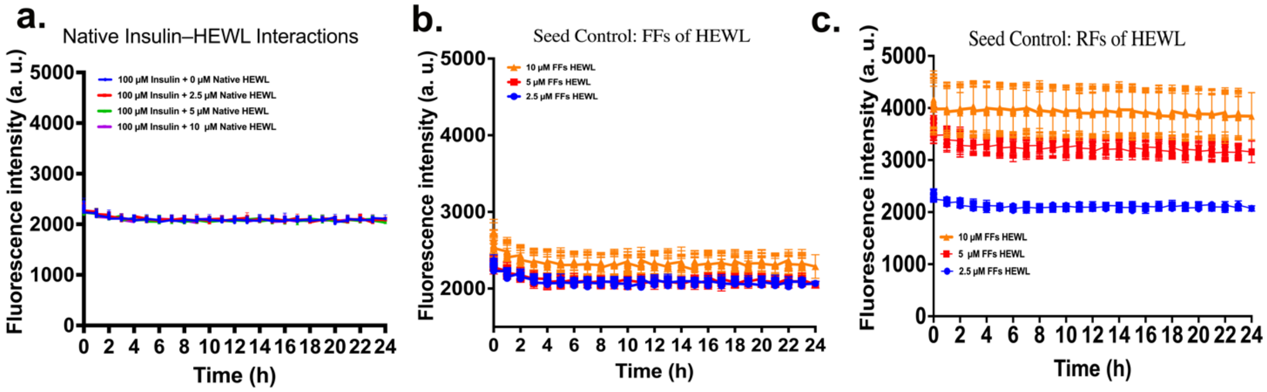


**Supplementary Figure S1. Thioflavin T fluorescence control experiments.** ThT fluorescence control experiments. (a) ThT fluorescence of native insulin (100 μM) incubated with increasing concentrations of monomeric HEWL (2.5, 5, and 10 μM) at pH 7.4 for 24 h. No significant increase in ThT fluorescence was observed, indicating that native insulin remains non-amyloidogenic under physiological conditions despite the presence of HEWL monomers. (b) ThT fluorescence of FFs HEWL seeds alone at 2.5, 5, and 10 μM. (c) ThT fluorescence of RFs HEWL seeds alone at the corresponding concentrations. The minimal and concentration-independent ThT signals from the HEWL seeds confirm that the enhanced ThT fluorescence observed in cross-seeding experiments originates predominantly from newly formed insulin amyloid fibrils rather than the seed material itself.
